# Unicycler: resolving bacterial genome assemblies from short and long sequencing reads

**DOI:** 10.1101/096412

**Authors:** Ryan R. Wick, Louise M. Judd, Claire L. Gorrie, Kathryn E. Holt

**Affiliations:** Centre for Systems Genomics, The University of Melbourne, Victoria, Australia; Department of Biochemistry and Molecular Biology, Bio21 Molecular Science and Biotechnology Institute, The University of Melbourne, Victoria, Australia

## Abstract

The Illumina DNA sequencing platform generates accurate but short reads, which can be used to produce accurate but fragmented genome assemblies. Pacific Biosciences and Oxford Nanopore Technologies DNA sequencing platforms generate long reads that can produce more complete genome assemblies, but the sequencing is more expensive and error prone. There is significant interest in combining data from these complementary sequencing technologies to generate more accurate “hybrid” assemblies. However, few tools exist that truly leverage the benefits of both types of data, namely the accuracy of short reads and the structural resolving power of long reads. Here we present Unicycler, a new tool for assembling bacterial genomes from a combination of short and long reads, which produces assemblies that are accurate, complete and cost-effective. Unicycler builds an initial assembly graph from short reads using the *de novo* assembler SPAdes and then simplifies the graph using information from short and long reads. Unicycler utilises a novel semi-global aligner, which is used to align long reads to the assembly graph. Tests on both synthetic and real reads show Unicycler can assemble larger contigs with fewer misassemblies than other hybrid assemblers, even when long read depth and accuracy are low. Unicycler is open source (GPLv3) and available at github.com/rrwick/Unicycler.

## 2 Introduction

Bacterial genomics is currently dominated by Illumina sequencing platforms. Illumina reads are accurate, have a low cost per base and have enabled widespread use of whole genome sequencing. However, much Illumina sequencing uses short fragments (500 bp or less) that are smaller than many repetitive elements in bacterial genomes^1^. This prevents short read assembly tools (assemblers) from resolving the full genome, and their assemblies are instead fragmented into dozens of contiguous sequences (contigs). Consequently, most available bacterial genomes are incomplete, which hinders large-scale comparative genomic studies.

Pacific Biosciences (PacBio) and Oxford Nanopore Technologies (ONT) sequencing platforms can sequence DNA fragments of 10 kbp or longer, but at a much higher cost per base than Illumina platforms. PacBio and ONT also have much higher per-base error rates than Illumina reads (5–15% vs <1%), although PacBio and ONT long reads are usually sufficient to complete bacterial genome assemblies with reasonable consensus accuracy^2,3^. Hence most researchers must choose between generating fragmented draft assemblies for many isolates with inexpensive Illumina sequencing, or generating complete assemblies for fewer isolates with expensive long read technologies. Hybrid assembly, which uses a combination of short and long reads, offers an alternative. In this approach, short reads are used to produce accurate contigs and long reads provide the information to scaffold them together. This requires relatively few long reads and can thus be the most cost-effective route to a complete bacterial genome.

Despite recent developments in long read technologies, Illumina reads are widely used in public health and research laboratories^4^, and are likely to remain so for some time due to their high accuracy and low cost. Moreover, Illumina data is already available for hundreds of thousands of bacterial isolates, and most of these are unlikely to be replaced with long read-only sequencing data. It is therefore likely that research and clinical labs will continue to use low cost Illumina reads for most samples and generate long reads as necessary to complete genomes of interest. Hybrid assembly, which requires fewer long reads than long read-only assembly, is the most cost-effective means of achieving this goal.

Multiple scaffolding tools exist to join Illumina contigs together using paired short reads or long reads, however mistakes are common and lead to structural errors (misassemblies) in the sequence^5^. Such errors could be avoided by performing scaffolding operations on assembly graphs – data structures which contain both contigs and their interconnections^6^ – rather than contig sequences alone. Here we present Unicycler, a new hybrid assembly pipeline for bacterial isolate genomes. Unicycler first assembles short reads into a highly accurate and connected assembly graph, and then scaffolds the graph with long reads. If the long read coverage is sufficient, it can produce a completed assembly with one contig per replicon. The assembly graph constrains possible paths through repeat regions, allowing Unicycler to achieve low misassembly rates. It is therefore an ideal assembler where the structural accuracy of the assembly is important.

## 3 Design and Implementation

To maximise ease of use, Unicycler encapsulates its entire pipeline (**Figure 1**) in a single command and automatically determines low-level parameters so users can expect good results with default settings.

**Figure 1:**
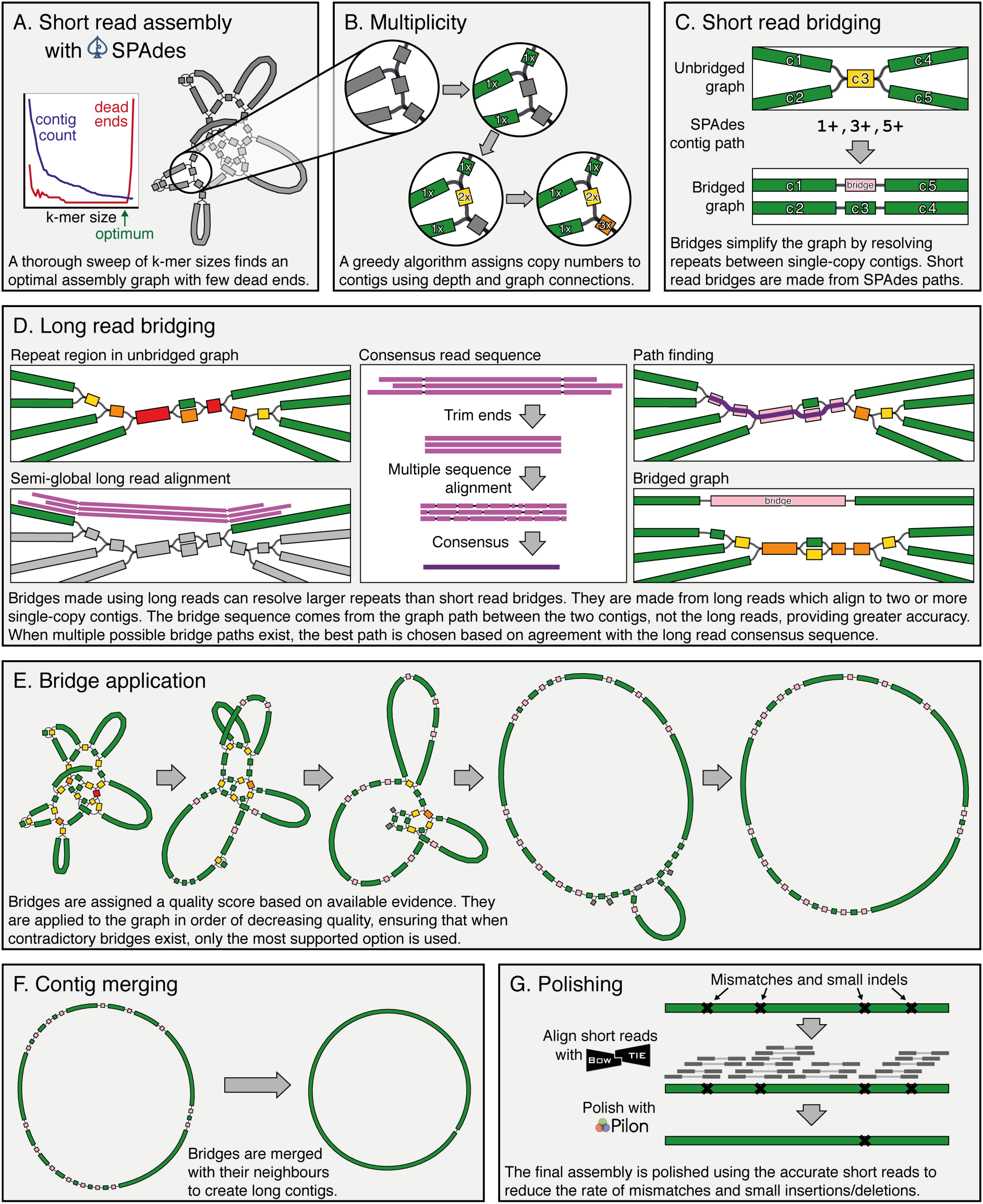
Key steps in the Unicycler pipeline.

### 3.1. Short read assembly

Unicycler uses SPAdes (v3.6.2 or later) to construct a De Bruijn graph assembly using a wide range of k-mer sizes: 10 values spanning 20–95% of the Illumina read length (not exceeding 127, the largest k-mer possible in SPAdes)^7^. In SPAdes, large k-mers result in more complete assemblies, but excessively large k-mers can cause a fragmented graph with dead ends. Unicycler assigns a score to each k-mer graph 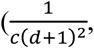 *c* = number of contigs, *d* = number of dead ends) and the highest scoring graph is selected as an ideal balance between minimising both contig count and dead ends (**Figure 1A**). As some contamination is possible in sequencing read sets (particularly when multiplexing, which is a common strategy for bacterial isolate sequencing on Illumina), Unicycler then removes contigs with a depth of less than half the median graph depth, unless doing so would create a dead end. This removes most contamination while leaving important graph structures intact.

### 3.2. Multiplicity

To resolve the graph as accurately as possible, Unicycler must distinguish between single-copy contigs (representing sequences that occur once in the genome, multiplicity *k* = 1) and repeat contigs (representing sequences that occur multiple times in the genome, multiplicity *k* > 1). When a bacterial genome consists of a single chromosome with no additional replicons, then for each contig *x*, its median read depth *d_x_* is a good indicator of its multiplicity *k_x_*. Single-copy contigs will have a median depth *d_x_* close to *D*, the median depth per base across the entire assembly, while repeat contigs will have a median depth near a multiple of that value (i.e. *d_x_* ~ *k_x_D*).

The relationship between median read depth and multiplicity is more complicated when the genome contains multiple replicons present at different copy number per cell. For example, small plasmids are often present in multiple copies, while large conjugative plasmids are often present in a single copy. To accommodate this, Unicycler also considers graph connections to help determine contig multiplicity. Repeat contigs typically have multiple graph connections at their start and end, while single-copy contigs usually have only a single connection at each end. These trends break down when the assembly graph is fragmented, which is one reason why Unicycler aims to minimise the number of dead ends when determining the optimal short read assembly graph. To determine multiplicity values, Unicycler therefore uses both depth and connectivity information. Initially, a multiplicity of one is assigned to all contigs that are near the median depth and have no more than one connection at either end. A greedy algorithm then propagates multiplicity where graph connections and depth are in close agreement (**Figure 1B**). This process is designed to distinguish single-copy contigs from repeats, even in plasmid-rich genomes.

### 3.3. Bridges

Unicycler scaffolds assembly graphs by constructing bridge contigs to connect pairs of single-copy contigs. Before bridging, single-copy contigs connect via multiple alternative paths containing one or more repeat contigs. After bridging, they connect via a simple, unambiguous path (**Figure 1C**). Bridges thus simplify the graph by resolving repeats. There are two primary sources of information available to Unicycler for creating bridges: read pair orientation and, more importantly, long reads.

### 3.4. Graph bridging using read pair information

When SPAdes assembles paired-end reads, it uses the read pair information to find paths through the assembly graph in a process known as repeat resolution (RR)^8^. SPAdes assemblies after RR (contigs.fasta output file) contain longer contigs than the assembly graph, but they are no longer in graph form. Unicycler uses SPAdes RR to build bridges and simplify the assembly while keeping it in graph form (**Figure 1C**). Furthermore, SPAdes RR may introduce misassemblies, so Unicycler excludes RR which spans a repeat longer than the typical read pair insert size, as such repeats are difficult to resolve with short read data alone. In this way, Unicycler produces an assembly graph which contains paired-end RR while avoiding some of the misassemblies that can occur in SPAdes.

### 3.5. Semi-global long read alignment

While short reads can resolve repeats up to the insert size of the library (typically <1000 bp), long reads provide a much more powerful source of scaffolding information. As a first step in long read bridging, Unicycler aligns all available long reads to the single-copy contigs. Since the long and short reads must be from the same biological sample, there should be no genuine structural discrepancies between the long reads and contigs. Semi-global alignment (end-gap free alignment) is therefore appropriate, where alignments can only clip when the end of a sequence is reached. Most available long read alignment tools such as BLASR^9^, BWA-MEM^10^, BLAST^11^ and LAST^12^ perform local alignment, so Unicycler implements semi-global alignment directly using the SeqAn C++ library^13^. To align low accuracy reads, it uses small alignment seeds (default of 10 bp). This results in many matches between a read and query, making a seed-extension alignment unfeasible. Instead, Unicycler searches for regions of high seed density and performs a banded alignment covering the best scoring regions.

### 3.6. Graph bridging using long read alignments

Long reads that align to multiple single-copy contigs can be used for bridging. Such reads contain a sample of the gap sequence between those contigs, and if multiple long reads connect a pair of contigs, Unicycler uses SeqAn to produce a consensus gap sequence^14,15^. Unicycler does not directly use this gap sequence in the bridge but instead uses it to find the best graph path connecting the contigs, via a branch and bound algorithm. Thus, the bridge sequence comes from the graph and reflects base calling accuracy of the short reads rather than the long reads that may have much lower accuracy (**Figure 1D**). Sometimes Unicycler cannot find a graph path connecting two single-copy contigs that are connected via long reads, such as when the short read graph is incomplete and contains dead ends. In these cases, the long read consensus sequence is directly used as the bridging sequence. Such bridges are more likely to contain errors, which is another reason why Unicycler strives to minimise dead ends in the assembly graph.

### 3.7. Bridge application

Having produced bridges from both short reads (SPAdes RR) and long reads, Unicycler can now apply them to simplify the graph structure (**Figure 1E**). Since some bridges may be erroneous, Unicycler assigns a quality score to each bridge and applies them in order of decreasingly quality, ensuring that when multiple contradictory bridges exist, the best-supported option is used. Bridge quality is a function of many factors: the number of reads which support the bridge (more is better); the alignment quality between the read consensus and graph path (higher identity is better); the length of the two contigs to be bridged (longer is better); the length and quality of the read alignments to the contigs (longer and higher identity is better); and the read depth consistency between the contigs (closer agreement is better). An ideal bridge connects two long contigs of the same depth, is supported by many reads with long alignments to the contigs, and has a graph path in close agreement with the read sequences.

### 3.8. Conservative, normal and bold

Unicycler can be run in three different modes that alter its cutoff for minimum acceptable bridge quality: conservative, normal and bold. In conservative mode, the quality cutoff is high (i.e. only very high quality bridges will be used). This mode is least likely to produce a complete assembly but carries a very low risk of misassembly and is appropriate for contexts where assembly accuracy is paramount. In bold mode, the quality cutoff is low (i.e. lower quality bridges will be used). This mode is most likely to complete the assembly but carries greater risk of error. It is suited to cases where completeness is more important than accuracy. Normal mode uses an intermediate cutoff and is appropriate in most cases.

### 3.9. Final steps

Following bridge application, Unicycler performs several actions to finalise the assembly graph. Contigs that have been used in bridges and no longer provide additional connection information are removed. Simple unbranching paths in the graph are merged to form long contigs (**Figure 1F**). Overlapping sequences at contig ends (created by the SPAdes assembly process) are removed so each contig’s sequence leads directly into its neighbours. If any circular replicon was completely assembled, it will now be a single contig with a link connecting its end to its start. A circular sequence can be shifted to any starting position without changing the biological information. Unicycler therefore uses TBLASTN to search for *dnaA* or *repA* alleles in each completed replicon^16^. If one is found, the sequence is rotated and/or flipped so that it begins with that gene encoded on the forward strand. This provides consistently oriented assemblies and reduces the risk that a gene will be split across the start and end of the sequence. As a final step, Unicycler uses Bowtie2 and Pilon to polish the assembly using short read alignments, reducing the rate of small errors (**Figure 1G**)^17,18^.

### 3.10. Included tools

Unicycler’s semi-global alignment algorithm is included as a stand-alone command line tool (unicycler_align), making it available for use in other pipelines. Unicycler also comes with a polishing tool (unicycler_polish) which applies variants identified by Pilon, GenomicConsensus^2^ and FreeBayes^19^, and assesses the assembly using ALE^20^. By iteratively polishing the genome with both short and long reads, this process can correct most errors in a completed assembly, including those in repeat regions.

## 4. Results

Unicycler’s performance was evaluated using read sets simulated for eight species, and using real read sets from the well-studied *E. coli* K-12 *substr*. MG1655. We further demonstrated the utility of Unicycler by using it to assemble the complete genomes of novel isolates of *Klebsiella pneumoniae* using newly generated Illumina, PacBio and ONT reads.

Unicycler’s performance was compared to that of alternative hybrid assemblers SPAdes (v3.8.1 in hybrid mode)^21^, ABySS (v1.9.0)^22^, npScarf (v1.6-01c)^23^ and Cerulean (v0.1)^24^. Default parameters and recommended settings were used for all tools (**Table S1**). The NaS tool can conduct hybrid assemblies^25^ but was excluded from this comparison because it depends on Newbler, a closed-source assembler only supported on RedHat/Fedora Linux. We also excluded ALLPATHS-LG^26^, which can perform hybrid assemblies but has strict library preparation requirements, restricting its applicability^27^.

### 4.1. Metrics

For both the simulated and real *E. coli* read tests, assemblies were evaluated by comparison to the corresponding complete reference genome using QUAST (v4.3)^28^. We focused on the following metrics: misassemblies, small error rate (mismatches and small indels) and NGA50.

QUAST identifies misassemblies as cases where a contig aligns to the reference genome in multiple pieces, not as a single continuous alignment, indicating a structural error in the contig. For the simulated read tests, reads were generated from the reference genome so misassemblies always indicate assembler mistakes. For the *E. coli* tests, there is not a perfect agreement between the reference genome and reads generated in different laboratories from different subcultures of *E. coli* K-12 *substr*. MG1655, so false positive misassemblies are possible.

The well-known N50 metric measures only contig size, not contig correctness, limiting its value. A large N50 can therefore result from inaccurately joining sequences into large misassembled contigs. By aligning contigs to a reference, QUAST produces more useful metrics including NGA50 (GA = ‘genome aligned’). Whereas N50 is based on contig lengths, NGA50 is based on the lengths of contig-to-reference alignments. A correctly assembled contig will have a single, unbroken alignment to the reference; a misassembled contig will be divided into multiple smaller alignments. Hence NGA50 is a metric for completeness that unlike N50 penalises misassemblies. In our tests, we used QUAST’s -- strict-NA option to break contigs at all misassembly locations, including local misassemblies, for particularly stringent NGA50 scores.

### 4.2. Simulated read sets

To provide a wide range of genome size and complexity, we simulated reads from 12 reference genomes from seven bacterial species (2 *Acinetobacter baumannii*^29,30^, *2 Escherichia coli*^31,32^ , 3 *Klebsiella pneumoniae*^33–35^, 1 *Mycobacterium tuberculosis*^36^, 1 *Shigella dysenteriae*^37^, 1 *Shigella sonnei*, 1 *Streptococcus suis*^38^) and the yeast *Saccharomyces cerevisiae*^39^ (**Table 1**). Plasmid and mitochondrial sequences were included at higher read depths, as appropriate.

**Table 1:** Reference genomes for simulated read sets.

| Species | Strain | Genome size (bp) | Description | Sequence type | Other features | GenBank assembly accession |
| --- | --- | --- | --- | --- | --- | --- |
| *Acinetobacter baumannii* | A1 | 3,917,739 | Circular chromosome, one plasmid | 231 | Large, repetitive biofilm-associated protein gene | GCA_000830055.1 |
| *Acinetobacter baumannii* | AB30 | 4,335,793 | Circular chromosome | 758 | Large, repetitive biofilm-associated protein gene | GCA_000746645.1 |
| *Escherichia coli* | K-12 MG1655 | 4,641,652 | Circular chromosome | 10 |  | GCA_000005845.2 |
| *Escherichia coli* | O25b:H4-ST131 EC958 | 5,249,449 | Circular chromosome, two plasmids | 131 |  | GCA_000285655.3 |
| *Klebsiella pneumoniae* | 30660/NJST258_1 | 5,540,936 | Circular chromosome, five plasmids | 258 |  | GCA_000598005.1 |
| *Klebsiella pneumoniae* | MGH 78578 | 5,694,894 | Circular chromosome, five plasmids | 38 |  | GCA_000016305.1 |
| *Klebsiella pneumoniae* | NTUH-K2044 | 5,472,672 | Circular chromosome, one plasmid | 23 |  | GCA_000009885.1 |
| *Mycobacterium tuberculosis* | H37Rv | 4,411,532 | Circular chromosome |  | High copy number PE and PPE genes | GCA_000195955.2 |
| *Saccharomyces cerevisiae* | S288c | 12,157,105 | 16 linear chromosomes, circular mitochondrial sequence |  | Eukaryote | GCA_000146045.2 |
| *Shigella dysenteriae* | Sd197 | 4,560,911 | Circular chromosome, two plasmids | 146 | High insertion sequence content | GCA_000012005.1 |
| *Shigella sonnei* | 53G | 5,220,473 | Circular chromosome, four plasmids | 152 | High insertion sequence content | GCA_000283715.1 |
| *Streptococcus suis* | BM407 | 2,170,808 | Circular chromosome, one plasmid | 1 |  | GCA_000026745.1 |

We used ART (v2.5.8) to generate synthetic paired-end reads from each reference genome with length (125 bp), insert size (400 bp) and error profile mimicking Illumina HiSeq 2500 reads^40,41^. Synthetic long reads were generated from each reference genome using PBSIM^42^ at three accuracies (60%, 75% and 90%), two mean lengths (10 kbp and 25 kbp) and eight depths (0x, 0.25x, 0.5x, 1.0x, 2.0x, 4.0x, 8.0x and 16.0x), yielding six short read sets and 42 hybrid read sets per strain. For the short read sets, we performed five assemblies: Unicycler in each of its modes (conservatives, normal and bold), SPAdes and ABySS. For the hybrid sets, we performed six assemblies: Unicycler (all modes), SPAdes, npScarf and Cerulean. All sets were generated in five replicates, resulting in 16920 total assemblies.

### 4.3. Assembly of simulated short read datasets

For assemblies of only synthetic short reads, Unicycler outperformed the other assemblers in each QUAST metric (**Figures 2** **and** **3**, **Table S2**). It is particularly interesting to compare Unicycler to SPAdes, since Unicycler uses SPAdes to build the initial short read assembly graph.

**Figure 2:**
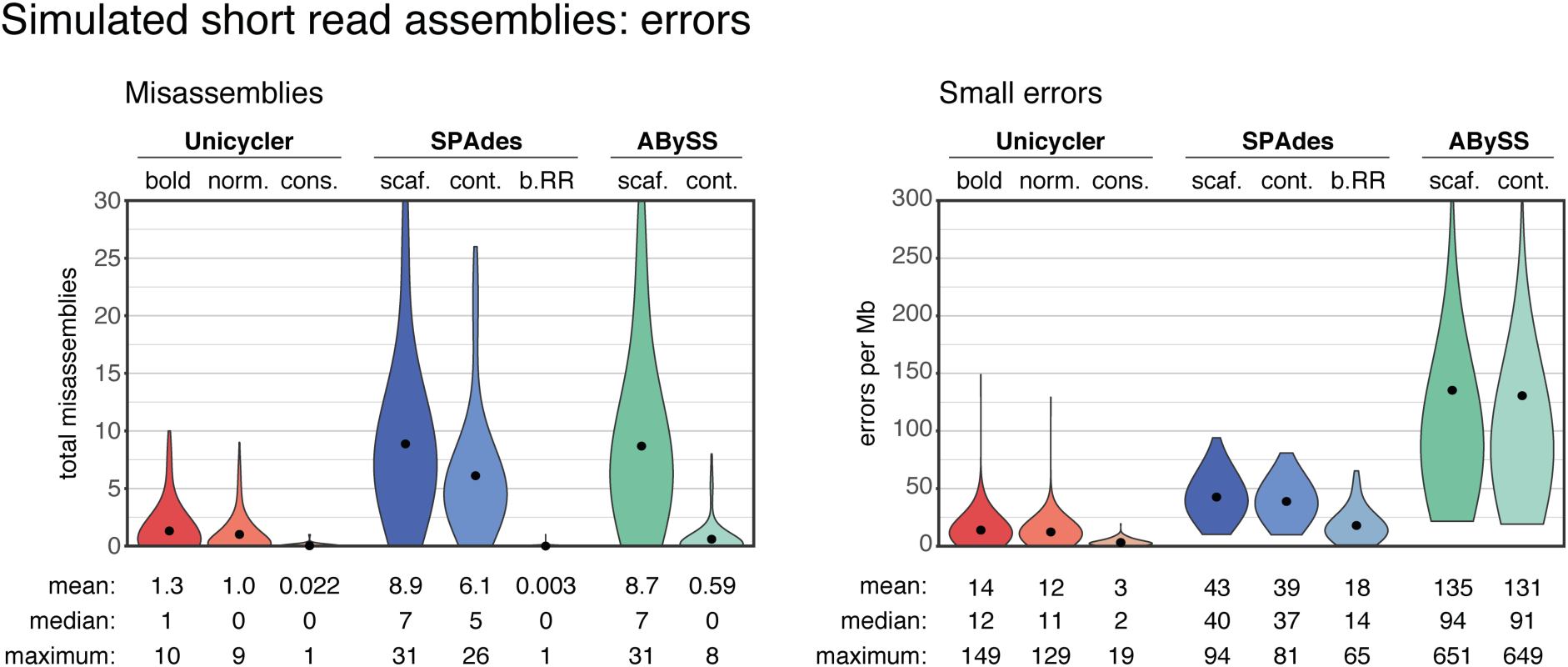
Misassembly and small error (mismatches and indels) rates for assemblies of simulated short read sets, summarising results across all reference genomes and replicate tests (total 360 per assembler).

**Figure 3:**
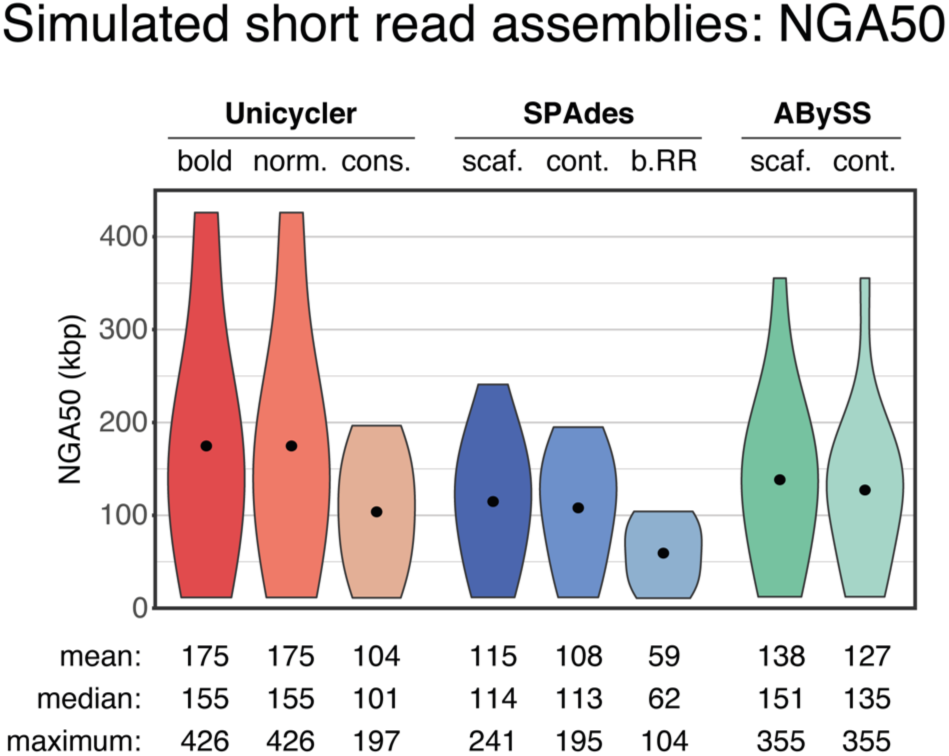
NGA50 for assemblies of simulated short read sets, summarising results across all reference genomes and replicate tests (total 360 per assembler).

In normal and bold modes, Unicycler achieved the most complete assemblies, as measured by NGA50. This is attributable to the wide k-mer range used in assembly. While SPAdes automatically selected a k-mer range of 21–55 and ABySS was run with a k-mer of 64 (the maximum value with default compilation settings), Unicycler typically chose 95 as its ideal k-mer, giving it greater power to assemble repetitive regions.

The misassembly rate of pre-RR SPAdes assemblies was very low, demonstrating that RR is the source of most misassemblies in SPAdes contigs. In conservative mode, Unicycler does not use SPAdes RR and therefore achieves similarly low misassembly rates. In normal and bold modes, Unicycler does use RR, but only if it exceeds a quality threshold. This selectiveness explains why normal and bold Unicycler assemblies have lower misassembly rates than the SPAdes contig assemblies from which they are derived.

### 4.4. Hybrid assembly of simulated long and short read datasets

Unicycler surpassed other assemblers when conducting hybrid assemblies of synthetic reads (**Table S2**). Misassembly rates in the hybrid assemblies were often much higher than in the short read only assemblies, illustrating the difficulty of resolving repeats with long reads (**Figure 4**). Both npScarf and Cerulean consistently produced assemblies with 10 or more misassemblies. SPAdes produced fewer misassemblies, but some genomes, particularly the *Shigella* genomes with many high copy number ~1 kbp repeats associated with insertion sequences, resulted in many errors. Unicycler’s misassembly rates were the lowest and correlated with the assembly mode (conservative, normal or bold).

**Figure 4:**
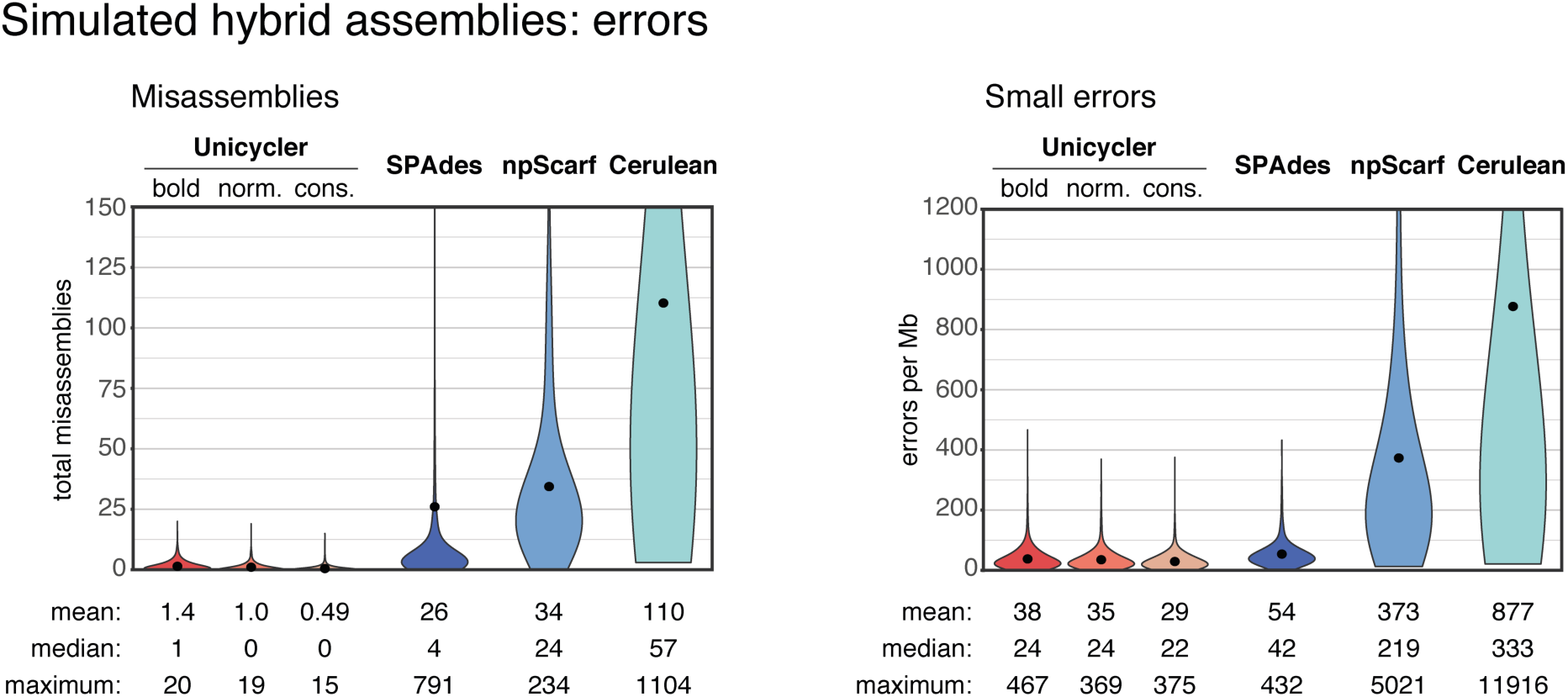
Error rates for hybrid assemblies of simulated short and long read sets, summarising results across all reference genomes and replicate tests (total 2520 per assembler).

Small error rates (mismatches and small indels) were lowest in Unicycler and SPAdes, as they both derive their final contigs from the short read assembly graph, not from the long read sequences. Unicycler and SPAdes both perform polishing, which may also contribute to their low small error rate. NGA50 was dependent on the long read depth, and Unicycler performed best at all tested depths (**Figure 5**). This is due to Unicycler’s low misassembly rates (other assemblers’ NGA50 scores were reduced due to their higher occurrence of misassemblies) and its ability to produce bridges using as few as one long read. In many cases, Unicycler produced complete or nearly complete assemblies with only 4x long read depth.

**Figure 5:**
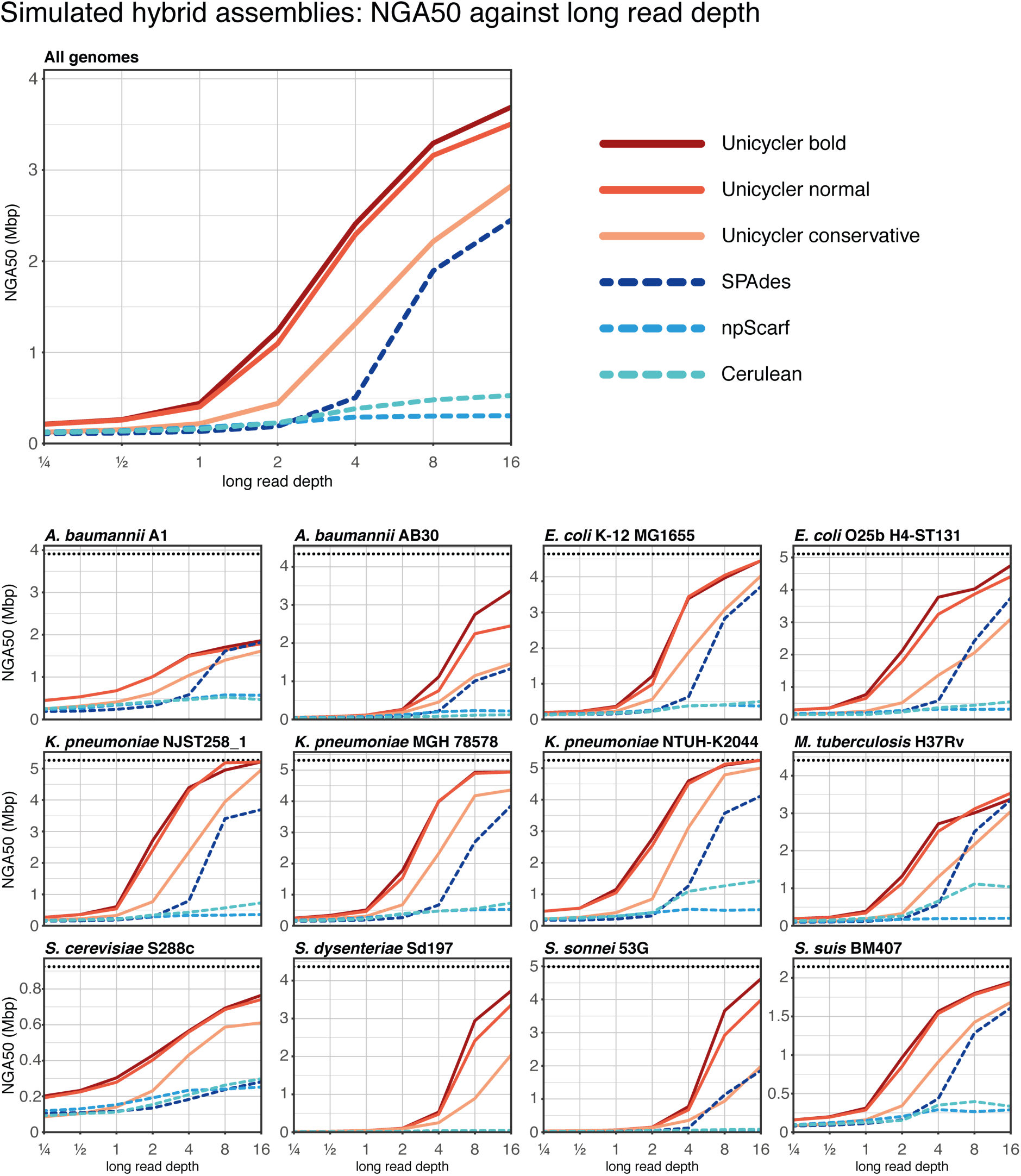
Mean NGA50 values for hybrid assemblies of simulated read sets. Mean values were calculated across all read lengths, read accuracies and replicate tests for each reference genome (210 hybrid read sets each); the top panel shows mean values for all 12 reference genomes (2520 hybrid read sets). Horizontal dashed lines indicate the N50 size of the reference genome. For the bacterial genomes, this is the size of their only chromosome; for *Saccharomyces*, it is the size of chromosome XIII, an intermediate-sized replicon in the genome.

Theoretical analyses of assembly show that error-prone reads are nearly as informative as error-free reads, suggesting that read accuracy is less important than length^43,44^. Unicycler’s performance on the simulated read sets matched these findings. Read length significantly affected the resulting NGA50 for Unicycler (all modes) and SPAdes (**Figure 6**). In contrast, read accuracy had a weaker effect on Unicycler’s NGA50 values, demonstrating its effectiveness in using long reads regardless of base calling accuracy.

**Figure 6:**
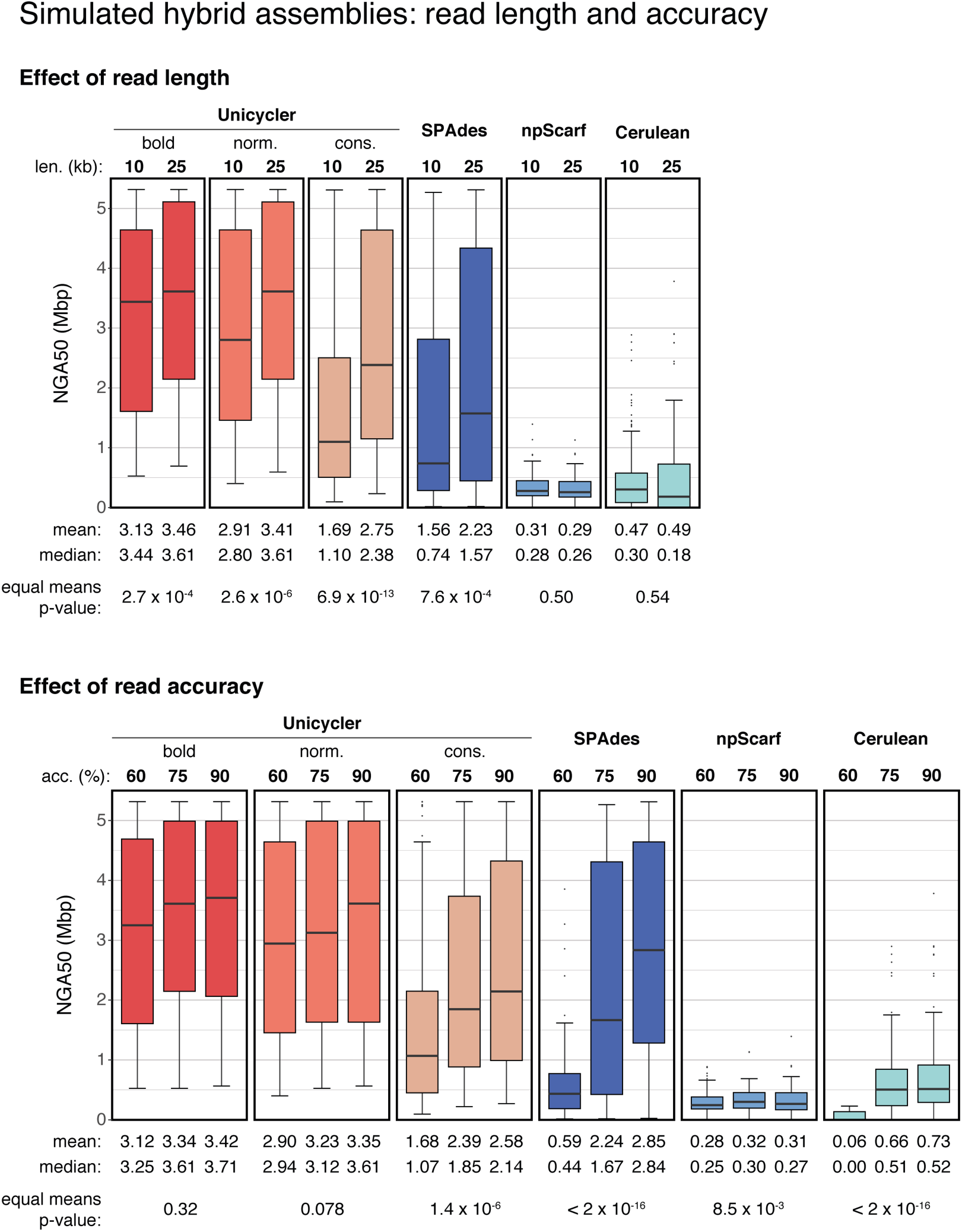
NGA50 values segregated by read length and read accuracy. These plots summarise results across all reference genomes and replicate tests, but only include the tests of 8x long read depth. For the read lengths, the p-value was made with a two-tailed t-test. For the read accuracies, the p-value was made with a one-way ANOVA test.

### 4.5. Computational performance

The assembly tests were all conducted with 8 CPU cores and 16 GB of RAM. Unicycler was slower than the alternative hybrid assemblers, taking a median time of 46 minutes to assemble the 8x long read depth synthetic tests. SPAdes and npScarf performed the fastest, both having a median time of 8 minutes and maximum time of less than 25 minutes on the same data. Cerulean had a median time of 23 minutes, although some Cerulean processes did not complete due to crashes or exceeding the 24-hour time limit. The complex biofilm-associated gene in *Acinetobacter baumannii* A1 was slow to bridge in Unicycler, resulting in a maximum run time of 13 hours. However, Unicycler did fully assemble this gene sequence in many read sets where the other assemblers produced a fragmented or misassembled result.

### 4.6. Real *E. coli* K-12 read sets

We tested the same assemblers using real reads of the *E. coli* K-12 *substr*. MG1655 genome. The short reads for these tests were produced by Illumina on their MiSeq platform. Long reads were from five different platforms: ONT R7, ONT R9, PacBio RS, PacBio RS II C2 chemistry and PacBio RS II C3 chemistry (**Table 2**). The ONT R9 reads were further split into two groups, pass and fail, as determined by ONT’s Metrichor software (v0.16.37960). For each platform and long read depth, we conducted 20 trials using different random subsamples of long reads at the same depths used for simulated data. Accuracy was assessed by comparison to the E. coli K-12 *substr*. MG1655 reference genome (accession NC_000913.3) generated using Sanger-based capillary sequencing at the University of Wisconsin in 1997^31^.

**Table 2:** Real *E. coli* K-12 *substr*. MG1655 read sets.

| Platform | Read count | Total length (Mbp) | Aligned length (Mbp) | Mean depth | Median length (bp) | N50 length (bp) | Mean identity (%) | Availability |
| --- | --- | --- | --- | --- | --- | --- | --- | --- |
| Illumina MiSeq | 2,200,000 | 584.7 | 581.7 | 125.3 | 300 | 300 | 99.4 | http://www.illumina.com/systems/miseq/scientific_data.html |
| ONT R7 | 68,455 | 311.5 | 151.4 | 32.6 | 3,751 | 7,290 | 67.7 | https://www.ncbi.nlm.nih.gov/sra/?term=ERR637419 |
| ONT R9 (fail) | 24,649 | 148.2 | 118.7 | 25.6 | 6,729 | 9,230 | 75.8 | http://lab.loman.net/2016/07/30/nanopore-r9-data-release/ |
| ONT R9 (pass) | 50,277 | 328.8 | 280.7 | 60.5 | 7,669 | 9,315 | 81.0 | http://lab.loman.net/2016/07/30/nanopore-r9-data-release/ |
| PacBio RS, C2 enzyme C2 chemistry | 42,582 | 106.8 | 105.1 | 22.6 | 2,016 | 4,030 | 85.0 | https://github.com/PacificBiosciences/DevNet/wiki/E-coli-K12-MG1655-Resequencing |
| PacBio RS II, P4 polymerase C2 chemistry | 82,590 | 437.0 | 432.7 | 93.2 | 4,589 | 7,566 | 80.7 | http://www.ncbi.nlm.nih.gov/sra/SRX669475 |
| PacBio RS II, P5 polymerase C3 chemistry | 47,910 | 396.8 | 395.7 | 85.3 | 7,613 | 11,789 | 85.0 | http://www.ncbi.nlm.nih.gov/sra/SRX533603 |

These tests showed the same trends as the simulated data: Unicycler achieved more complete assemblies at lower long read depths than other assemblers (**Figure 7**, **Table S3**). Notably, Unicycler performed worst on the set with shortest reads (PacBio RS) not the set with lowest identity (ONT R7), while it performed best on the set with longest reads (PacBio RS II C3). This matches the simulated results and the theoretical predictions, again showing that read length is more important than accuracy for Unicycler.

**Figure 7:**
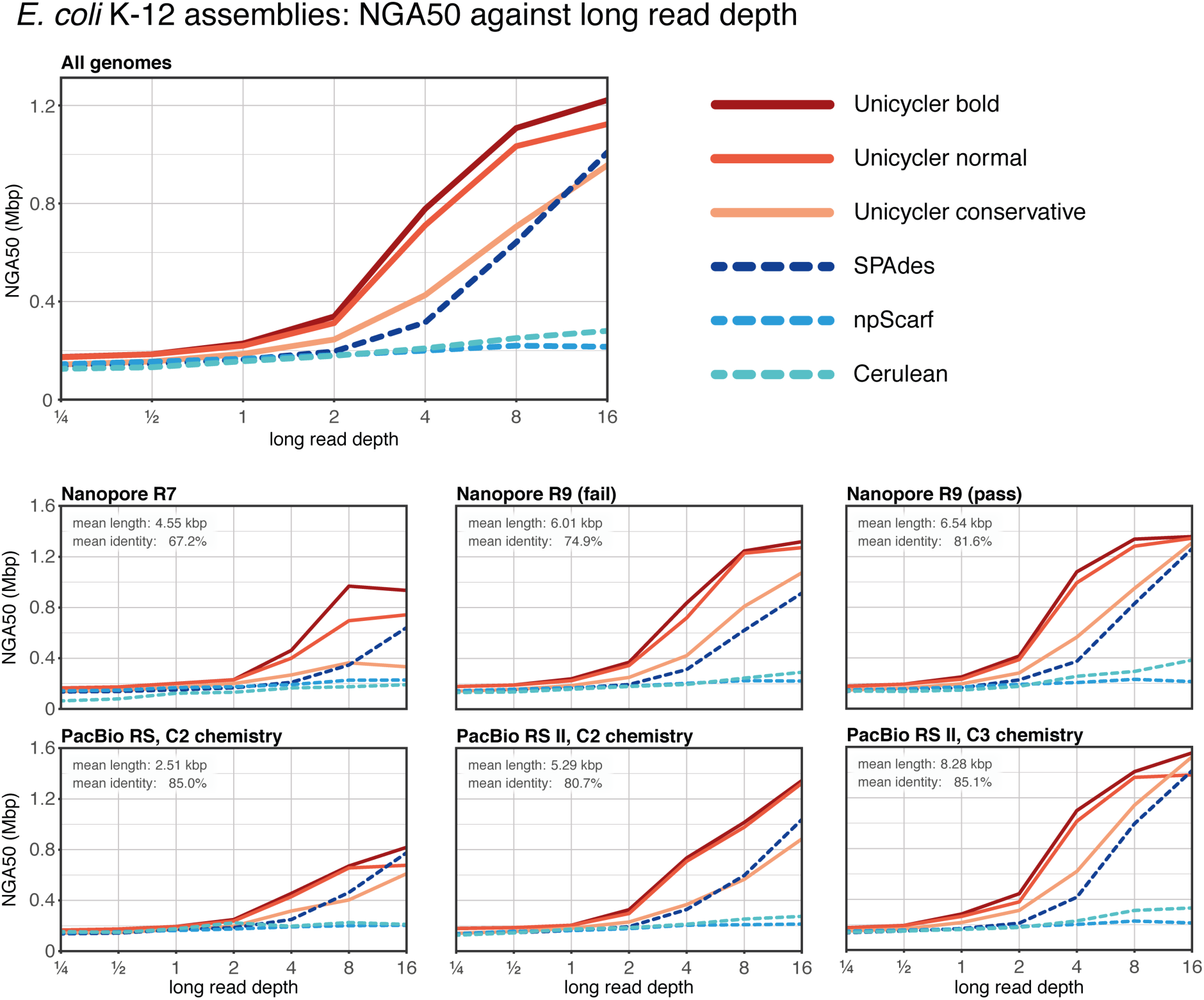
Mean NGA50 values for hybrid assemblies of real E. coli read sets, summarised across 20 replicate tests at each depth. Top panel shows mean values for all six long read sets.

The NGA50s for these tests were markedly lower than those obtained with reads simulated from the *E. coli* K-12 *substr*. MG1655 reference genome. With simulated reads, both Unicycler and SPAdes were often able to achieve complete or near-complete assemblies. With real reads, Unicycler and SPAdes best NGA50 values were 2.0 Mbp and 1.4 Mbp, respectively. This appears to be due to false positive misassembly calls resulting from genuine differences between the reference genome sequence and the genomes of the isolates that were sequenced using Illumina, PacBio and ONT platforms. For example, one copy of IS*l*A in the Illumina read set relocated to a position 105 kbp away from the position in the reference genome; when an assembly spanned this region, QUAST identified the difference as a misassembly, reducing the NGA50.

### 4.7. *Klebsiella pneumoniae* PacBio assembly case study

To explore the utility of Unicycler, we used it to assemble the genome of a *K. pneumoniae* isolate, INF274, which was difficult to assemble using alternative techniques. INF274 is a multidrug resistant sequence type (ST) 340 strain isolated from the urine of a Melbourne hospital patient who had a urinary tract infection. It belongs to clonal group (CG) 258, a common cause of multidrug resistant hospital-associated infections globally. This isolate was first sequenced on Illumina HiSeq 2500 (generating 750 Mbp of 125 bp paired-end reads) and then on a PacBio RS II (generating 1.3 Gbp of long reads, many of which exceeded 10 kbp in length). Both reads sets are high quality and are an ideal set of inputs for hybrid assembly. Notably, the library preparation for the PacBio reads followed a standard size-selection protocol that excluded short fragments of DNA, so small plasmids were underrepresented in the long reads.

We performed hybrid assemblies using Unicycler (normal mode) and SPAdes (v3.8.1), and PacBio-only assemblies using HGAP (v3) and Canu (v1.3). Since this is a novel isolate, we did not analyse the assembly results with QUAST. Instead, we compared the assemblies to each other and analysed the alignment of Illumina reads to each (**Figures 8** **and** **S2**).

**Figure 8:**
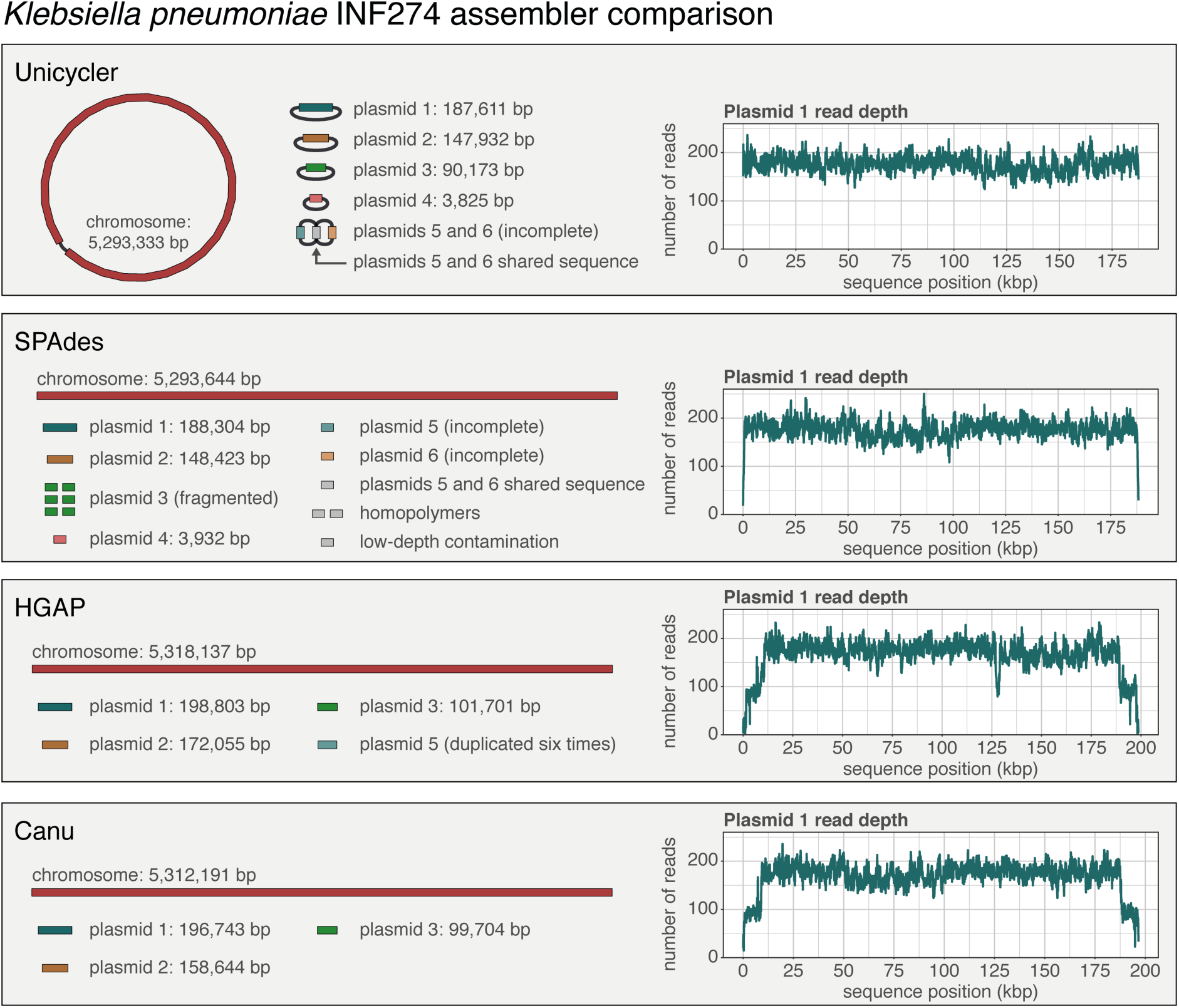
Final assemblies of Klebsiella pneumoniae INF274 produced by Unicycler, SPAdes, HGAP and Canu. The contigs/graph of the assembly are shown on the left, coloured by replicon. The read depth plot of plasmid 1 is shown on the right. Low depth at the ends of the contig is indicative of overlapping ends.

Of the four assemblers, only Unicycler and Canu produce a graph file for their final assembly, but Canu did not circularise any replicons, so the sequences remained linear. When viewed in Bandage^6^, the Unicycler graph clearly distinguished between replicons that formed completed circularised sequences and those that did not. Only plasmids 5 and 6 remained incomplete in the Unicycler assembly, as they contain shared sequence and there was a lack of PacBio reads for these replicons, preventing Unicycler from scaffolding them apart. SPAdes and HGAP output their assemblies only as linear sequences, making it difficult to make the distinction between complete and incomplete replicons. The SPAdes assembly suffered from the same problem as Unicycler with plasmids 5 and 6, and it failed to assemble plasmid 3, which contains a prophage sequence also present in the chromosome.

Since Unicycler’s graph-based scaffolding naturally results in circular sequences, it did not have duplicated sequences at the start/end of circular replicons. SPAdes suffered from a slight overlap and both HGAP and Canu had significant overlaps, indicated by the drop in read depth near the ends of contigs. These must be repaired manually or with a tool such as Circlator^45^.

### 4.8. *Klebsiella pneumoniae* ONT assembly case study

To assess Unicycler’s performance on low-depth ONT datasets, we performed R9 sequencing on *K. pneumoniae* isolate INF125, a virulent ST45 strain isolated from the urine of a Melbourne hospital patient. Reads were generated over a four-hour period resulting in a total of 156 Mbp of sequence (depth=28.6x) with an N50 length of 11,470 bp. ONT streams sequence data as it is generated, making it technically feasible to analyse the data in real time and stop sequencing when a complete assembly is reached. To investigate the suitability of the assemblers for this task, we assessed their performance over time by generating 240 subsets of reads, one set per minute of sequencing, each containing all reads generated up to that minute (e.g. set 60 contained all reads generated in the first hour of sequencing). All sets were assembled with three hybrid assemblers (Unicycler normal mode, hybridSPAdes and npScarf) and one long read-only assembler (miniasm^46^). We assessed each assembly using N50, number of contigs, and error rates when aligning the Illumina reads to the assembly (**Figure 9**, **Table S4**). A high read alignment identity is indicative of a low small error rate (mismatches and small indels). A high proportion of concordantly aligned reads is indicative of a low misassembly rate.

**Figure 9:**
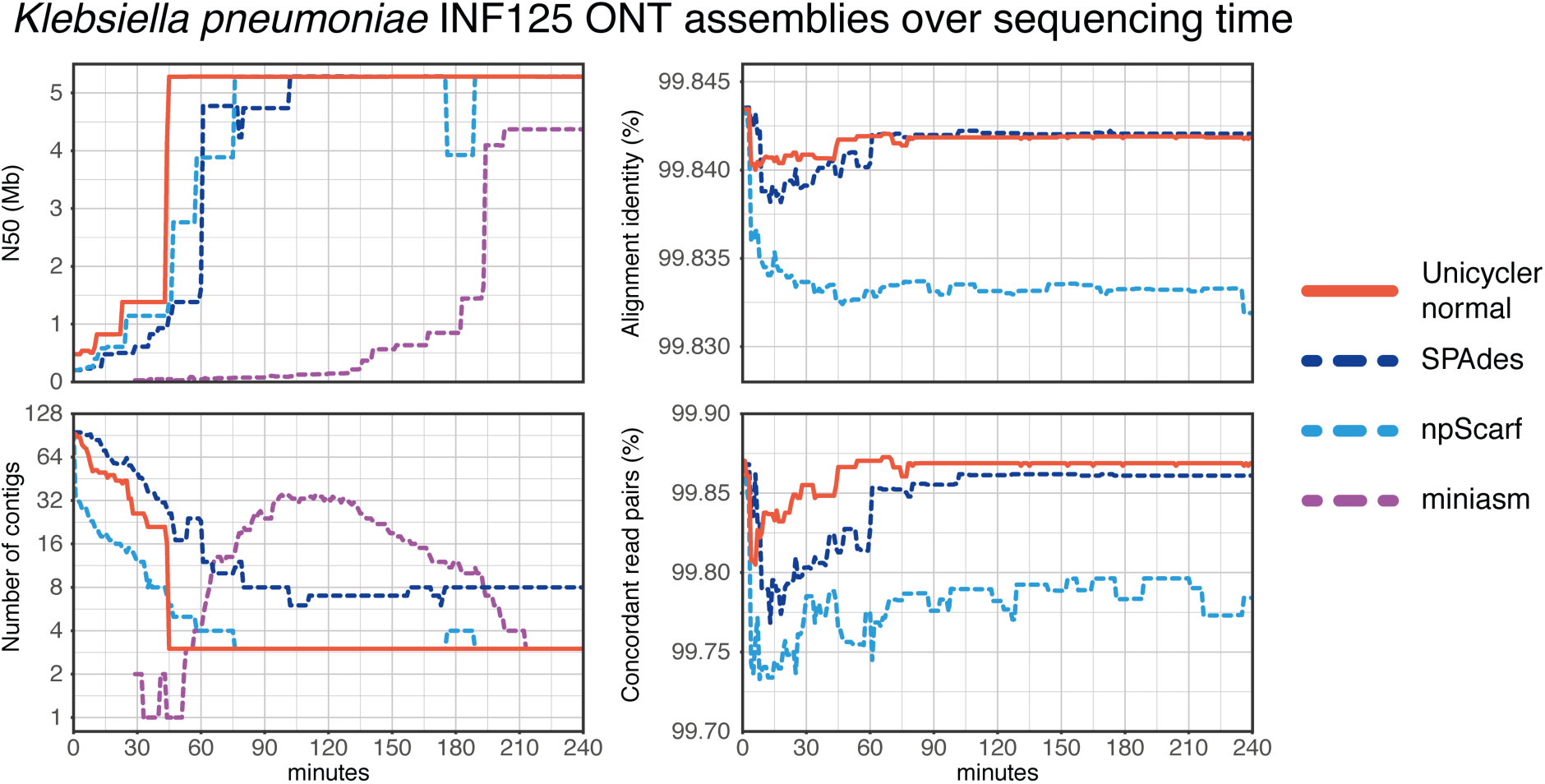
Assembly metrics of Klebsiella pneumoniae INF125 produced by Unicycler, SPAdes, npScarf and miniasm over a four-hour period of sequencing. Miniasm assemblies contain error rates comparable to that of the raw reads and is therefore excluded from the error rate plots.

Unicycler was the first assembler to complete the assembly at 45 minutes (depth=5.3x). npScarf completed the assembly at 76 minutes (9.0x), SPAdes took 102 minutes (12.1x) and miniasm, which uses only long reads, required 213 minutes (25.3x). All completed assemblies contained three contigs (one chromosome and two plasmids), except for SPAdes assemblies that contained extra, erroneous contigs. Read error metrics show that both Unicycler and SPAdes consistently produced more accurate assemblies than npScarf, although the magnitude of the benefit was small (**Figure 9**).

### 4.9. Limitations

As Unicycler operates on a short read assembly graph, it requires high quality short reads. Specifically, it is important that there are very few unsequenced regions of the genome that create dead ends in the assembly graph. Unicycler’s performance will suffer if the assembly graph is fragmented and incomplete.

### 4.10. Summary

Unicycler performed well on both short read-only data sets and all types of hybrid read sets, producing more complete assemblies than other assemblers. Perhaps more importantly, Unicycler produced fewer misassemblies than other assemblers, which often had unacceptably high error rates. As long read sequencing becomes more common, so will completed genome assemblies, enabling new research into genome structure. High quality assemblies free of structural errors, such as those produced by Unicycler, will be critical to research in this field.

## 5. Availability and Future Directions

Unicycler’s primary use case is when a researcher wishes to complete the assembly of an isolate for which Illumina reads already exist. To facilitate this, future development of Unicycler will add streaming support for ONT sequencing, using reads to create and update bridges in the graph in real time during a sequencing run. This will allow users to halt sequencing once a genome is sufficiently resolved, conserving sequencing resources for other isolates. This modality is currently possible with npScarf, however in our tests Unicycler was more accurate than npScarf and reached complete assemblies with lower read depths. Future development will also focus on improving Unicycler’s computational performance. Unicycler is not currently able to perform large assemblies such as human genomes and metagenomes. Algorithmic improvements to long read alignment, path finding and graph manipulations will all be required for Unicycler to be appropriate in such cases.

Unicycler is open source (GPLv3) and available at github.com/rrwick/Unicycler.

## 7. Supplementary data

**Figure S1:**
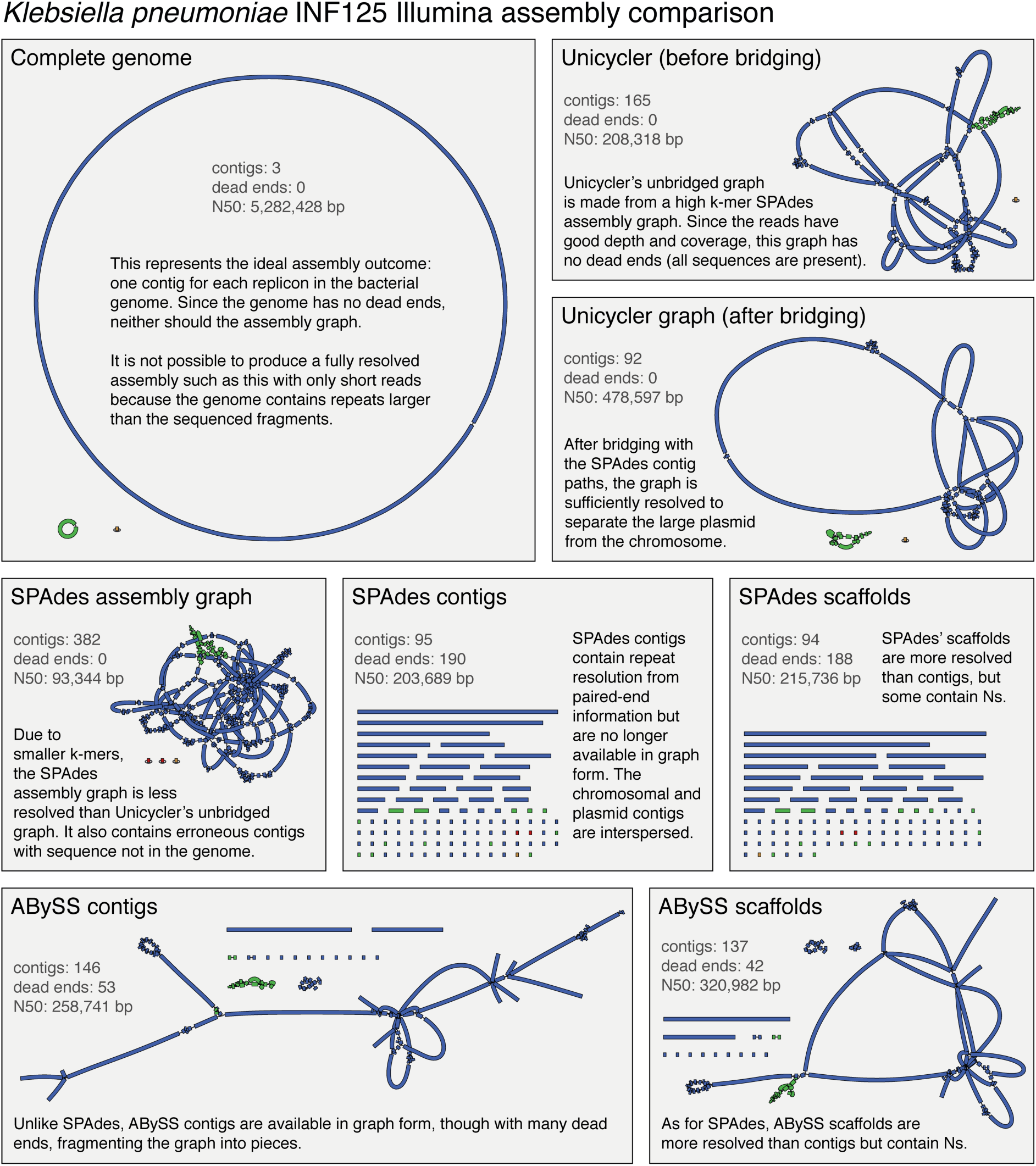
Comparison of assembly graphs made by different assemblers using only short reads for *Klebsiella pneumoniae* INF125. Contigs are coloured based on which replicon they represent.

**Figure S2:**
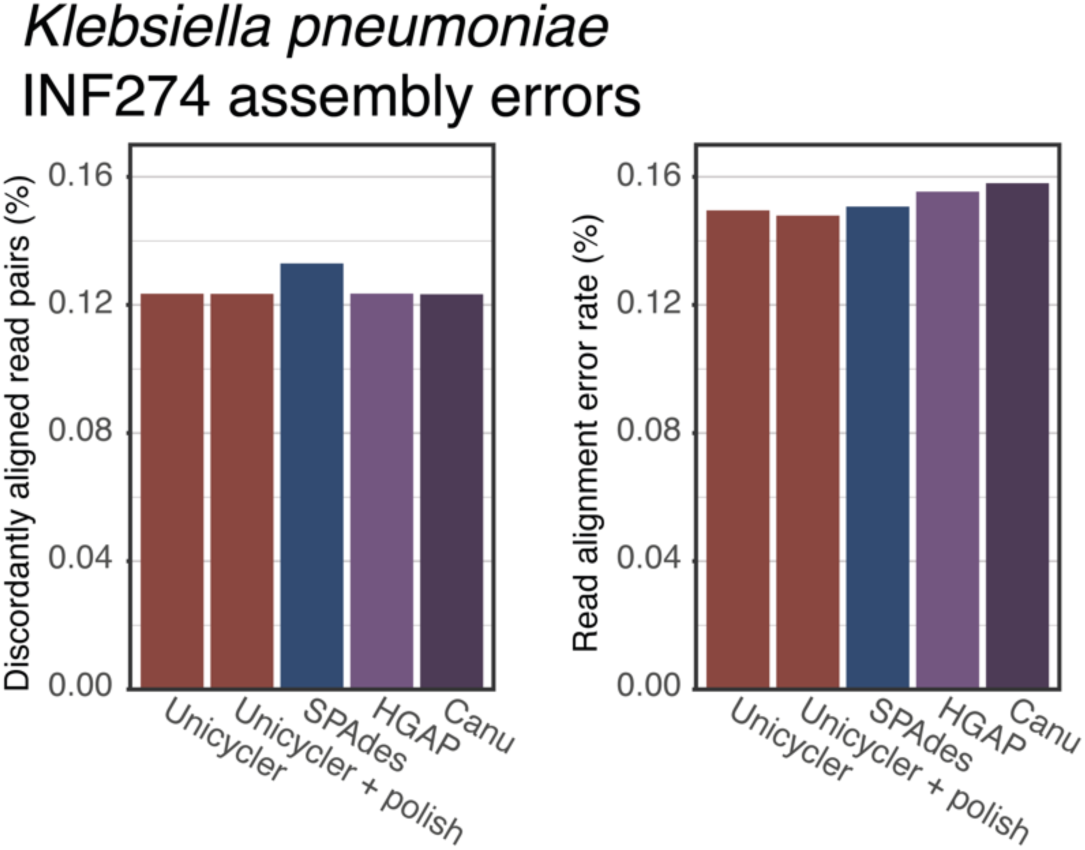
Error rates when aligning Illumina reads to each *Klebsiella pneumoniae* INF274 assembly. An increase in discordant pairs is indicative of a misassembly. An increase in error rate is indicative of small errors (mismatches and small indels).

**Table S1: Commands used for assembly, evaluation and read simulation.**

**Table S2: Raw QUAST results for simulated read tests.**

**Table S3: Raw QUAST results for *E. coli* read tests.**

**Table S4: Raw quality metrics for *K. pneumoniae* INF125 ONT assemblies over time.**

